# Concerted genome expansion of heritable symbionts in an insect host

**DOI:** 10.1101/2025.05.06.652349

**Authors:** Emily A Hornett, Masayuki Hayashi, Keisuke Nagamine, Steve Paterson, Daisuke Kageyama, Gregory D D Hurst

**Affiliations:** Department of Evolution, Ecology & Behaviour, University of Liverpool, Liverpool, UK; Department of Vector Biology, Liverpool School of Tropical Medicine, Liverpool, UK; Institute for Plant Protection, National Agriculture and Food Research Organization, Kannondai 2-1- 8, Tsukuba, Ibaraki 305-8666, Japan; Institute of Agrobiological Sciences, National Agriculture and Food Research Organization (NARO), 1-2 Owashi, Tsukuba, Ibaraki 305-8634, Japan

**Keywords:** *Spiroplasma*, *Rickettsia*, symbiont, genome size, mobile genetic element, prophage, insertion sequence

## Abstract

Maternally transmitted symbionts represent important components of arthropod biology, acting as both beneficial partners and as reproductive parasites. Their strict vertical transmission greatly reduces their effective population size, making selection less efficient and driving a pattern of molecular evolution where genome size reduces. Contrastingly, there are sporadic observations of genome expansion in symbionts from clades where the genome has previously reduced in size. It is currently unclear whether these events are an idiosyncratic consequence of exposure to novel active mobile elements, or are driven by the host context. In this paper, we report the concerted genome expansion of two co-infecting heritable symbionts that supports a host-related driver of expansion. We assembled the genomes of *Spiroplasma* and *Rickettsia* bacteria that co-infect the lacewing *Mallada desjardinsi*. Both symbionts had the largest genome reported to date in their clade, approximately twice the median size for their respective genus. Genome expansion was driven by proliferation of mobile elements in both cases, but the underpinning elements were distinct. The observation that the proximate causes of expansion differed between symbionts led us to reject the hypothesis that concerted expansion was driven by a common mobile element invasion. We hypothesize these processes are driven either by bottleneck events impacting host population size, by the host environment causing stress-induced activation of mobile genetic elements, or by both symbionts having undergone a recent and coincident transition to vertical transmission.

## Introduction

Microbial genomes are highly compact in comparison to those of eukaryotes, characterized by high coding density with little intergenic material. This compactness is largely a product of the high efficacy of selection consequent on the very large population size that microbes attain^1^. In addition, whilst bacteria can carry mobile genetic elements (MGEs) such as insertion sequences (IS) and prophage^2^, these do not commonly dominate the genomic constitution in the manner observed in many eukaryotes, where a significant fraction of the genome is often comprised of active or relic transposable elements^3^.

The evolutionary landscape of vertically transmitted microbial symbionts is distinct to other microbes^4^. Heritable microbes reside within eukaryotic, particularly arthropod, hosts and are most commonly transmitted maternally from a female host to her progeny within or on the outside of eggs. With a tight bottleneck at the point of vertical transmission, the effective population size for these microbes becomes aligned to the number of infected female hosts, leading to a reduced efficacy of purifying selection against mildly deleterious mutations. Heritable symbionts thus represent microbes that have core molecular evolution properties aligned to their eukaryotic hosts^4^.

One contrast with their eukaryotic host is that the decline in effective population size in heritable symbionts drives a well evidenced evolutionary pathway to genome shrinkage^4^. Weakened purifying selection in eukaryotes is commonly associated with increased genome size arising from replication and spread of transposable elements fuelled by recombination and sexual reproduction. Contrarily, whilst the initial evolution of heritable symbiosis may involve IS element and/or prophage accumulation^5,6,7^, the mutational bias to deletion in asexual heritable microbes drives genome shrinkage and streamlining^8^. This process has resulted in the smallest bacterial genomes observed naturally, for example *Nasuia deltocephalinicola*, an obligate endosymbiont of the Aster Leafhopper, with a genome size of just 112 Kb^9^.

Notwithstanding this general pathway, however, there are increasing reports where particular symbiont lineages have shown genome expansion events. This process was first observed in a member of the Rickettsiales, *Orientia tsutsugamushi*, whose 2.0-2.5 Mb genome was considerably larger than other members of the bacterial order (typically 1-1.5 Mb). Here, the accumulation of MGEs and accessory genes such as transposases, *tra* genes, phage integrases and reverse transcriptases, and proliferation of an integrative and conjugative element called ‘Rickettsiales amplified genetic element’ (RAGE) led to repetitive elements comprising almost 50% of the genome^10,11^. A further study revealed the particularly large genome of another rickettsial symbiont with its ∼1.8 Mb genome comprising ∼35% RAGEs alongside transposases and other MGEs^12^. More modest genome expansions have also been observed for other bacteria such as *Wolbachia* and *Amoebophilus asiaticus*^13,14,15^.

Heritable symbiont genome evolution thus presents contrasts, with weakened purifying selection and deletion bias driving reduced genome size, but then occasional lineages evidencing genome expansion through mobile element proliferation. An unresolved issue is the circumstances that lead to the occasional genome expansion events observed. A null hypothesis view is that it is random with respect to the lineages where this occurs – perhaps consequent on stochastic exposure of a mobile element that can then proliferate. The alternate hypothesis is that there are particular circumstances that enable proliferation. One prediction of the alternate hypothesis of host-driven factors is that heritable symbionts that co-infect a host would be predicted to have correlated evolutionary trajectory. Put simply, if the host provides the context driving genome expansion, it should be seen in both co-infecting symbionts.

Here, we provide a case study that supports the prediction that the host environment can drive heritable symbiont genome expansion. We sequenced and assembled the genome of the lacewing *Mallada desjardinsi*, to aid understanding of the interaction between the lacewing and two of its symbionts. The first symbiont is a member of the bacterial genus *Spiroplasma*, which includes insect pathogens, insect-vectored plant pathogens, and heritable symbionts of insects^16^. The strain in *Mallada desjardinsi* is heritable and expresses a male-killing phenotype where infected male hosts are exclusively killed during embryogenesis or early larval stages^17^. The second symbiont present is a member of the bacterial genus *Rickettsia*, a group of obligate host-associated intracellular microbes whose members are arthropod symbionts or arthropod-vectored pathogens^18^. The impact of *Rickettsia* on the lacewing host is not known, aside not being associated with sex ratio distortion. We report the symbiont genome sizes, analysed the cause of the genome expansions observed, resolved the relatedness of these strains to other members of their genus and compared their genomes to previously sequenced congeneric strains.

## Materials and Methods

### DNA isolation and long-read sequencing

A single female *Mallada desjardinsi* lacewing was selected from a 6^th^ generation inbred line originating from Matsudo, Japan. This line was co-infected with two strains of bacteria: a *Rickettsia* and a *Spiroplasma*. To prepare the lacewing tissue for DNA isolation, the whole adult insect was first snap frozen in liquid nitrogen and the tissue ground immediately using a sterile pestle. HMW DNA was then isolated using a Qiagen Genomic-tip 20/G and associated buffers following the manufacturers protocol. The resultant DNA was sequenced on a PacBio Sequell II SMRT Cell (HiFi) in CCS run mode. Reads were pre-processed using PacBio SMRT Link (version 10.2.0.133434) for adaptor removal and removal of CCS reads with a quality score (Q) <20. This resulted in a total of 1003881 reads, summing 12148926671 bases, with 98.4% being >Q20 and 96.4% being >Q30.

### Metagenome assembly and symbiont genome retrieval

The genomes of the lacewing host plus the two bacterial symbionts were assembled as a metagenome using Hifiasm^19^ (version 0.19.8-r603), with haplotig duplication purging. This metagenome assembly then underwent three rounds of polishing with the PacBio reads using Racon^20^ (version 1.5.0). The resulting draft metagenome, comprising lacewing, *Spiroplasma* and *Rickettsia* genomes, consisted of 233 contigs, with a total length of 470 Mb, and a N50 of 71 Kb. Contigs were taxonomically assigned to the lacewing, *Spiroplasma* or *Rickettsia* using a combination of Blobtools2^21^ (version 3.0.0), minimap2^22^ (version 2.24-r1122) and BLAST+^23^ (version 2.12.0+). The *Spiroplasma* and *Rickettsia* metagenome assembled contigs (MAGs) were separately assessed for genome size, number of contigs, completeness and contamination using QUAST^24^ (version 5.0.2,), BUSCO^25^ (version 5.2.2) and CheckM2^26^ (version 0.1.3) (Table S1).

### Genome size comparison

For Mollicutes, to which *Spiroplasma* belongs, the size in bp of sequenced complete genomes were collated from the database Molligen4 (https://services.cbib.u-bordeaux.fr/molligen4). For the Rickettsiales, to which *Rickettsia* belongs, data was extracted from Castelli *et al*.^27^. The genome size of a selection of these (107 Mollicutes, and 139 Rickettsiales) were plotted using R^28^ (version 3.6.3) with the packages ggplot2^29^ (version 3.5.0), dplyr^30^ (version 1.1.4) and forcats^31^ (version 1.0.0), alongside the symbionts reported here, to visualise the size of focal symbionts in the context of their taxonomic groups.

### Symbiont genome annotation

Coding sequences (CDS) of the main chromosome for both *Rickettsia* and *Spiroplasma* were identified and annotated using BAKTA^32^ (version 1.9.3). The *Rickettsia* and *Spiroplasma* main chromosomes were also functionally annotated using anvi’o^33^ (version 8), running the commands anvi-run-kegg-kofams and then anvi-estimate-metabolism to estimate KEGG pathway completeness. DFAST^34^ (version 1.3.0) was employed to ascertain the number of pseudogenes in each genome. Putative plasmid sequences were identified as being circular extra-chromosomal contigs with coverage higher than the main chromosome.

### Mobile genetic element and repeat detection

The main chromosomes of the *Rickettsia* and *Spiroplasma* symbionts carried by the lacewing host, along with genomes of congeneric representatives were analysed for the presence of repeat sequences and common mobile genetic elements. For *Rickettsia*, 32 additional genomes were included (genome accessions in Table S5), while for *Spiroplasma*, 54 additional genomes were included (genome accessions in Table S6). Only genomes where the main chromosome was complete (one contig) were used in the comparative study. The overall repetitive content of the genomes was assessed and visualised using Mummer4^35^ (version 4.0.0rc1). Prophage regions, Insertion Sequences (IS) and Tandom Repeats (TRs) were identified using PHASTEST^36^ (https://phastest.ca), ISEScan^37^ (version 1.7.2.3) and Tandom Repeat Finder^38^ (version 4.09.1) respectively. The presence of CRISPR repeats and spacers were investigated using CRISPR-CasFinder^39^ with default parameters (https://crisprcas.i2bc.paris-saclay.fr/CrisprCasFinder/Index). The results from these analyses were plotted using R^28^ (version 3.6.3) with the packages ggplot2^29^ (version 3.5.0), dplyr^30^ (version 1.1.4) and forcats^31^ (version 1.0.0).

For the *Rickettsia* symbiont, the main chromosome was additionally analysed for the presence of RAGEs as these elements have previously been shown to be important in *Rickettsia* genome expansion events^11,10,12^. For this a search was made for the presence of a definitive sequence of *tra* and associated genes, as described in Giengkam *et al*.^10^.

### Phylogenomic analyses

The phylogenetic relationship of the symbionts under study were analysed alongside selected representatives from their respective genera - the same strains as was included in the comparative analysis (Tables S5-S6). The phylogenies for both *Rickettsia* and *Spiroplasma* were based upon sequence data of 67 single copy core orthologs identified as being present in all genomes, then concatenated and aligned in anvi’o^33^ (version 8). The best protein model was identified using ModelFinder^40^ and a Maximum Likelihood tree was reconstructed in IQTree2^41^ (version 2.3.0), and visualised using FigTree (version 1.4.4, https://github.com/rambaut/figtree).

According to Bayesian Information Criterion, for the *Rickettsia* phylogeny, the best-fit model chosen was Q.plant+F+R4, and for the *Spiroplasma* phylogeny, the best fit model was LG+F+I+R5. *Orientia tsutsugamushi* and *Mycoplasmoides genitalium* were used as outgroups for the *Rickettsia* and *Spiroplasma* trees, respectively.

To ascertain whether particular IS elements were shared between the symbiont genomes, the phylogenetic relationships of transposase sequences from the shared IS families were estimated. For this, where a particular IS family was identified as being found in both bacterial genomes, the longest transposase coding sequence was chosen from each bacterium, and from selected representatives from the *Spiroplasma* and *Rickettsia* genera. For each IS family, the amino acid sequences were aligned using MEGAX^42^ (version 11.0.13). The best protein model was identified using ModelFinder^40^ and a Maximum Likelihood tree was reconstructed in IQTree2^41^ (version 2.3.0), and visualised using FigTree (version 1.4.4, https://github.com/rambaut/figtree).

## Results

### The *Rickettsia* and *Spiroplasma* main chromosomes are the largest sequenced within their respective clades

The *Spiroplasma* and *Rickettsia* genomes retrieved from the lacewing metagenome were each assembled into one complete circular bacterial chromosome, plus 3 circular plasmids for *Rickettsia*. The presence and number of *Spiroplasma* plasmids remains unresolved due to copious repetitive sequences that make their status ambiguous. For *Spiroplasma*, hereafter named strain *s*Md, the main chromosome is 3137804 bp, and for *Rickettsia*, hereafter named strain *r*Md, the main chromosome is 3245895 bp. CheckM2 and BUSCO analyses indicated the genomes were 99-100% and 96-100% complete for core marker sets respectively (Table S1). Both genomes are to our knowledge the largest complete genomes sequenced to date in their respective genera. Moreover, *Rickettsia r*Md is the largest sequenced complete genome within the order Rickettsiales (Figure S1), and the *Spiroplasma s*Md is the largest sequenced complete genome within the class Mollicutes (Figure S2).

### The *Spiroplasma & Rickettsia* genomes are highly repetitive

In order to investigate the factors driving the size expansion of both of the symbiont main chromosomes, we analysed whether the genomes contained repeat sequences. While the BUSCO results for both genomes demonstrate that the core genes themselves are not for the most part duplicated (1/151 and 3/364 genes for *Spiroplasma* and *Rickettsia*, respectively; Table S1), the main chromosomes both show very high levels of repetition (Figure 1). Observation of the plots reveal a difference in the repeat pattern between the main chromosomes of *Rickettsia* and *Spiroplasma. Rickettsia* is dominated by repeats 200-5000 bp in length, while *Spiroplasma* has a larger number of repeats in the >5 Kb range than *Rickettsia*.

**Figure 1.**
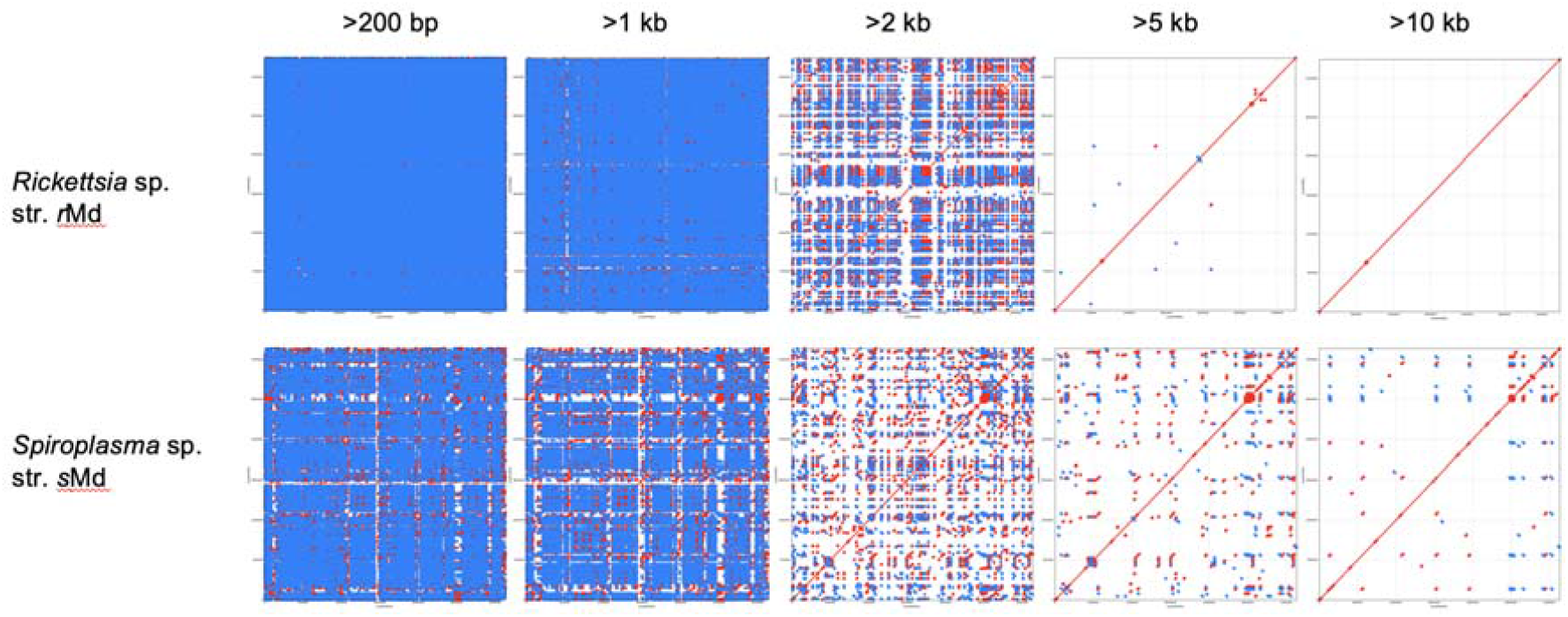
Mummer plots depicting repeat sequences within the *Rickettsia r*Md and *Spiroplasma s*Md main chromosomes. Repeats were filtered by size then plotted. For each graph, the X-axis and the Y-axis is the same; i.e. the main chromosome is plotted against itself. Repeats are shown in blue (forward direction), and red (reverse direction).

To ascertain the level of diversity in repetitiveness across *Rickettsia* and *Spiroplasma*, repetitive sequences in the main chromosome of four representatives from each genus were similarly visualised using mummerplot and compared to the strains isolated from the lacewing (Figures S3 and S4). Our focal symbionts displayed a broader signal of repetition than other members of the genus.

### The *Rickettsia* and *Spiroplasma* genomes are dominated by insertion sequences (IS) and/or prophage

Annotation of the *Rickettsia r*Md main chromosome revealed that of 3818 coding sequences (CDS), almost one third of these (1114) were classed as transposases or IS accessory proteins (Table S2). When analysing the genome for full IS, which typically comprise a transposase flanked by short repeat sequences, the *Rickettsia* main chromosome carried 1564 IS totalling 1778924 bp and comprised 54.81% of the main chromosome. In addition to IS, *Rickettsia* also had 142 tandem repeats, and one intact phage region of 41.4 Kb. While a sequence of *tra* genes in the *Rickettsia* genome indicate the presence of a putative RAGE, it is unclear whether this is functional given interruption by numerous transposases. There was no evidence of a CRISPR-Cas system (Figure 2A).

**Figure 2.**
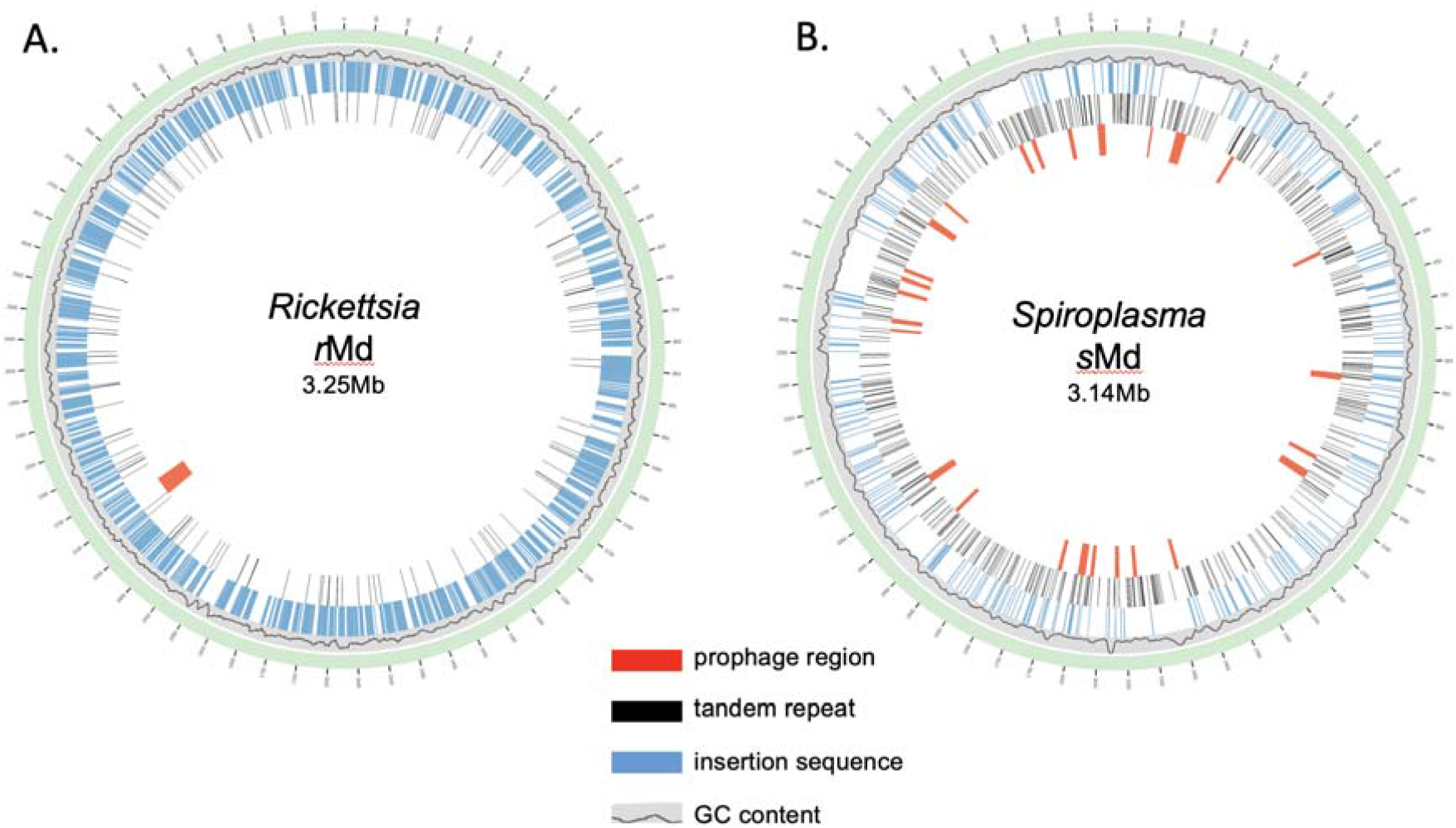
Circos plots depicting GC content and mobile genetic element features of the **A**. *Rickettsia r*Md and **B**. *Spiroplasma s*Md main chromosomes.

The *Spiroplasma s*Md main chromosome has 4430 CDS, with 407 being transposases (just under 10% of the CDS; Table S3). There were 309 full IS comprising transposase plus flanking repeats, totalling 432487 bp. In addition, of the CDS annotated, 1177 were assigned as being phage related (contain key words: phage, virus, capsid, integrase, terminase, baseplate), making up ∼25% of the CDS (Table S3). When the presence of prophage within the *Spiroplasma* main chromosome was investigated further, 26 phage regions were discovered, totalling 276.1 Kb. Further repetitive sequences were identified in the form of 621 tandem repeats. Again, there was no evidence of a CRISPR-Cas system. (Figure 2B - circos plot).

### *Rickettsia* and *Spiroplasma* do not share IS

Given the presence of large numbers of IS in both the *Rickettsia* and *Spiroplasma* main chromosomes, we investigated whether the particular IS found were shared between the two bacterial strains, reflecting lateral gene transfer as a driving factor of the concerted genome expansions. The IS situated in the main chromosome of *Rickettsia* belong to 17 different IS families (Table S4). Four of these IS families are shared with those found in *Spiroplasma* (IS3, IS5, IS30, IS481) - with these four being the only IS families present in *Spiroplasma*. However, the frequencies of the IS differ between the two symbionts. For *Spiroplasma*, by far the most prevalent IS are from family IS30 (n=269), fewer from IS481 (n=33), and IS3 and IS5 are rare (n=3 and n=4, respectively). In contrast, of these four IS families the reverse is true; in *Rickettsia* IS3 is the most prevalent (n=506), then IS5 (n=23), then IS481 (n=21), with IS30 being the least prevalent (n=7).

For both *Spiroplasma* and *Rickettsia*, transposase coding sequence from each of the four IS families were aligned alongside sequence from selected congeneric strains, and four phylogenies estimated. For each IS family, the transposase sequences cluster with those from their respective genera rather than with each other (Figures S5-S8). Taken together, the results indicate the IS elements are not shared by *Spiroplasma* and *Rickettsia* co-infecting the lacewing host by direct transfer from one symbiont to the other.

### *Rickettsia* and *Spiroplasma* genomes retain a high level of functionality

There was a high level of pseudogenisation in *Rickettsia r*Md and a moderate level in *Spiroplasma s*Md as ascertained by DFAST (Rickettsia 1179 pseudogenes, 926 of which are transposases; Spiroplasma 286 pseudogenes, 40 of which are transposases). However, KEGG analysis reveals that both genomes retain a high level of functionality, with module completeness being similar to other members of their respective genera (Figures S9-13).

### IS elements and prophage regions vary in presence and frequency across *Rickettsia* and *Spiroplasma*

To investigate whether there is a general relationship between main chromosome size and the presence of IS and prophage, a selection of sequenced complete genomes (main chromosome in one contig) were analysed and compared to that of the *Rickettsia* strain *r*Md isolated from the lacewing *M. desjardinsi* (Figure 3, Table S5). *Rickettsia r*Md lies within the bellii group of *Rickettsia*, which comprises both vertically inherited and pathogenic bacteria. As previously mentioned, this *Rickettsia* has the largest genome in the genus, and also the highest number of IS elements. Tellingly, the second largest *Rickettsia* strain, isolated from the spider *Oedothorax gibbosus* at 2.62 Mb^43^, has the second largest number of IS, indicating that proliferation of IS also drove genome expansion in this bacterium. While the lacewing *Rickettsia* does have one prophage, it is not a driving factor in its genome expansion, and other *Rickettsia* within the genera have higher numbers of prophage regions.

**Figure 3.**
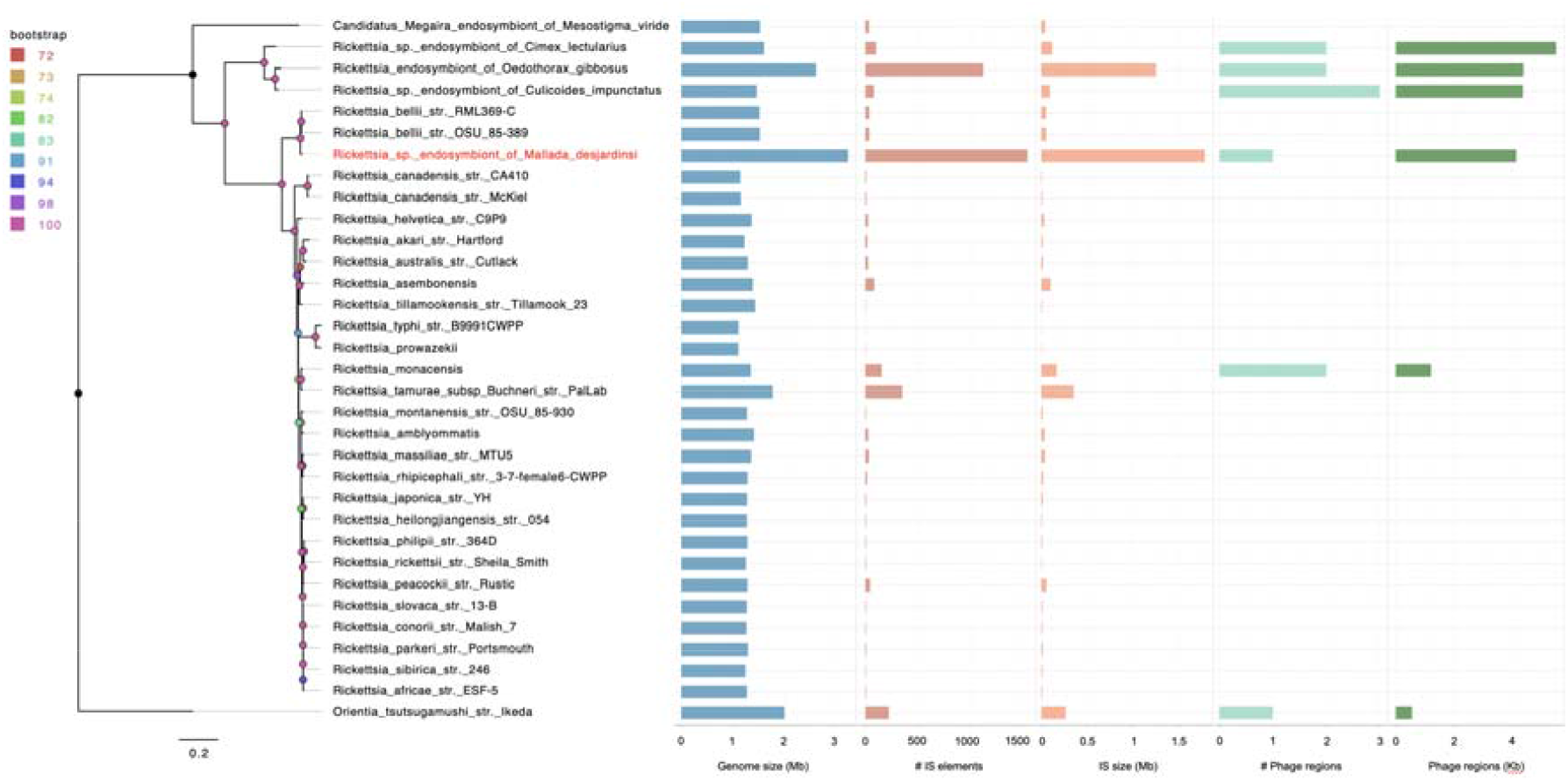
Phylogeny of *r*Md (red) and selected *Rickettsia* strains with sequenced complete genomes. Genome size (Mb), the number of IS elements along with the total size of IS (Mb), and the number of phage regions with total size (Kb) is also given alongside.

Similarly, a selection of sequenced complete *Spiroplasma* genomes (main chromosome in one contig) were analysed and compared to that of the *Spiroplasma* strain *s*Md isolated from the lacewing *M. desjardinsi* (Figure 4; Table S6).

**Figure 4.**
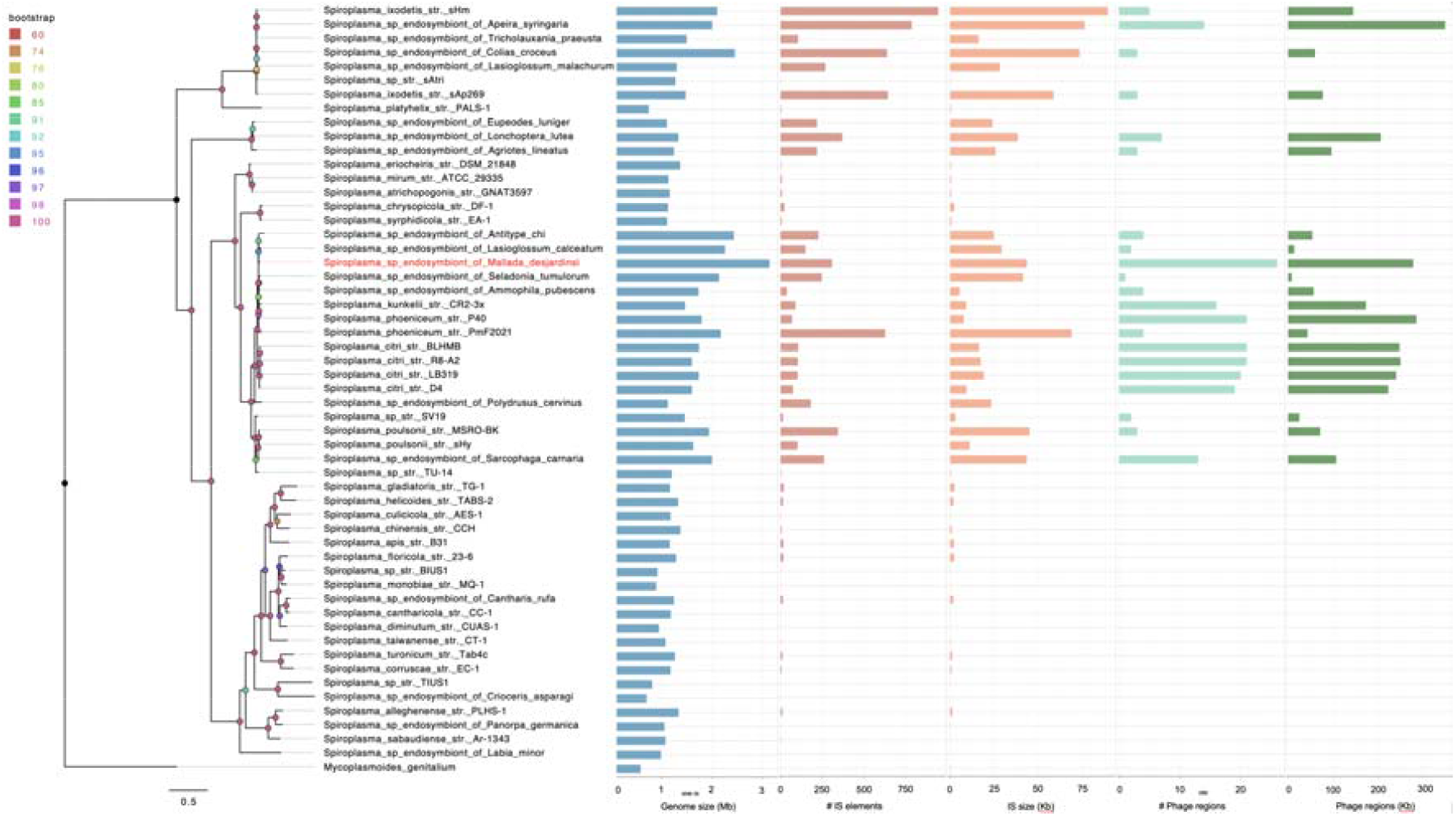
Phylogeny of *s*Md (red) and selected *Spiroplasma* strains with sequenced complete genomes. Genome size (Mb), the number of IS elements along with the total size of IS (Mb), and the number of phage regions with total size (Kb) is also given alongside.

*Spiroplasma s*Md is most closely related to infectiously transmitted insect vectored plant pathogens including *S. citri* and *S. phoeniceum*. As previously stated, *s*Md has the largest main chromosome sequenced as a complete genome to date. While it does have numerous IS, others within the genus have comparable or higher numbers of IS, and while proliferation of IS may be a contributing factor driving genome expansion in these strains, it is a contributor alongside other repetitive elements. *Spiroplasma s*Md does however, have the highest number of prophage regions, although the total size is not the highest observed in the genus. Looking across the *Spiroplasma* strains, it appears that a combination of proliferation of IS and prophage leads to genome expansion within this genus.

## Discussion

The pattern of molecular evolution of bacteria alters when they become heritable, moving from large effective population sizes to ones paralleling their eukaryotic host. The widely observed genome streamlining that results from alignment, however, contrasts with the eukaryotic host – the bias to deletion mutations in microbial genomes combines with the weakened efficacy of purifying selection to result in progressive pseudogenization and ultimately smaller genomes of very high coding content^8^. Accumulation of mobile elements following initial evolution to host association has been observed previously^7^, as a phase prior to genome streamlining. In our study, mobile genetic element proliferation occurs in the context of a previous history of genome shrinkage within the genera.

The presence of overarching drivers of genome expansion is supported by our observation of expansion occurring in co-infecting symbionts. The lacewing *Mallada desjardinsi* carries two symbionts (that can co-exist within individual lacewings) both showing genome expansion, and both having the largest recorded main chromosome in their group (in the case of *Rickettsia*, the wider Rickettsiacae, in the case of *Spiroplasma*, Mollicutes). The *Rickettsia* genome included three large plasmids that are also IS rich, two of which are themselves larger in size than the previously observed largest plasmid recorded in the *Rickettsia* (448 Kb & 172 Kb vs *pacificus* at 121 Kb^44^).

The proximate causes of genome expansion in the two microbes were distinct – *Spiroplasma* genome expansion was largely driven by prophage accumulation, *Rickettsia* by diverse IS elements. Moreover, the IS-mediated genome expansion of *Rickettsia r*Md is distinct from the genome expansion of *Orientia* and other *Rickettsia*, in being associated with a single mobile element type (rather than combinations of IS elements and RAGE). Overall, genome expansion is a convergent phenotype occurring in heritable symbionts associated with diverse repetitive elements.

The concerted expansion we observed supports a currently as yet uncharacterized driver of genome expansion, rather than genome expansion occurring idiosyncratically without biological drivers. We can exclude a common exposure to horizontally transmitted elements – the intracellular arena scenario^45^ – as the genome expansion events have a different main core cause (IS vs prophage). The involvement of different elements in expansion also indicate expansion is not particular to the type of element. There are three non-mutually exclusive explanations to explain the concerted expansion. First, the host provides a common environment in terms of effective population size, driving expansion by host population size restriction. Second, the host provides a shared physiological environment. This environment could include stress sensed by the bacteria, which can impact both IS and phage activity^46,47^. Third, expansion of symbiont genomes through prophage and IS element proliferation have occurred following initial transition of the microbe from free-living/infectious to host association^5,7^. Contrary to this hypothesis, both our symbionts derive from clades that are already host associated^18,48^. However, the strains may have both transitioned recently to vertical transmission. The *Spiroplasma* in *M. desjardinsi* is closely allied to arthropod-vectored plant pathogens e.g. *S. phoeniceum*^49^. The bellii clade of *Rickettsia* contains both heritable symbionts and vectored pathogens^50,51^.

One observation from our work is that heritable microbes can tolerate these high loads of mobile elements, which in the case of the *Rickettsia* comprise around half the genome and a doubling in size compared to most members of the genus. KEGG analysis indicated no obvious degradation of symbiont metabolic function compared to non-expanded relatives. Thus, purifying selection remained sufficiently strong to maintain the function of core genes. Heritable microbes experience both within and between host selection, and it is expected that mutations causing loss of core functions would compromise within host competitiveness and thus be eliminated. For the *Spiroplasma*, the novel prophage material may have both costs and benefits. Prophage commonly carry a cargo of symbiosis-relevant genes, as observed in the *Arsenophonus* genome expansion^5^.

In conclusion, the evolution of vertical transmission is associated with reduced effective population size. Whilst the most common consequence of this reduction is genome streamlining, sporadic cases of genome expansion through mobile element activity are observed, paralleling the processes that lead to genome expansion in eukaryotic host taxa. Our observation of high levels of expansion in two co-infecting symbionts, with both recorded as the largest in their bacterial groups, indicates a core underpinning cause of expansion. However, that this expansion is caused by distinct repetitive elements indicates that a host factor is the driving force rather than the nature of the mobile element itself.

## Acknowledgements

This work was supported by a collaborative grant from the UKRI and JSPS to GH, SP and DK (NE/S012346/1).

## References

1. Lynch M, Conery JS. 2003 The Origins of Genome Complexity. Science 302, 1401–1404. (doi:10.1126/science.1089370)

2. Lang AS, Buchan A, Burrus V. 2025 Interactions and evolutionary relationships among bacterial mobile genetic elements. Nat Rev Microbiol, 1–16. (doi:10.1038/s41579-025-01157-y)

3. Wells JN, Feschotte C. 2020 A Field Guide to Eukaryotic Transposable Elements. Annu Rev Genet 54, 539–561. (doi:10.1146/annurev-genet-040620-022145)

4. McCutcheon JP, Boyd BM, Dale C. 2019 The Life of an Insect Endosymbiont from the Cradle to the Grave. Current Biology 29, R485–R495. (doi:10.1016/j.cub.2019.03.032)

5. Siozios S et al. 2024 Genome dynamics across the evolutionary transition to endosymbiosis. Current Biology 34, 5659-5670.e7. (doi:10.1016/j.cub.2024.10.044)

6. Moran NA, Plague GR. 2004 Genomic changes following host restriction in bacteria. Current Opinion in Genetics & Development 14, 627–633. (doi:10.1016/j.gde.2004.09.003)

7. Plague GR, Dunbar HE, Tran PL, Moran NA. 2008 Extensive Proliferation of Transposable Elements in Heritable Bacterial Symbionts. J Bacteriol 190, 777–779. (doi:10.1128/JB.01082-07)

8. Mira A, Ochman H, Moran NA. 2001 Deletional bias and the evolution of bacterial genomes. Trends in Genetics 17, 589–596. (doi:10.1016/S0168-9525(01)02447-7)

9. Bennett GM, Moran NA. 2013 Small, Smaller, Smallest: The Origins and Evolution of Ancient Dual Symbioses in a Phloem-Feeding Insect. Genome Biol Evol 5, 1675–1688. (doi:10.1093/gbe/evt118)

10. Giengkam S, Kullapanich C, Wongsantichon J, Adcox HE, Gillespie JJ, Salje J. 2023 Orientia tsutsugamushi: comprehensive analysis of the mobilome of a highly fragmented and repetitive genome reveals the capacity for ongoing lateral gene transfer in an obligate intracellular bacterium. mSphere 8, e00268–23. (doi:10.1128/msphere.00268-23)

11. Cho N-H et al. 2007 The Orientia tsutsugamushi genome reveals massive proliferation of conjugative type IV secretion system and host–cell interaction genes. Proc. Natl. Acad. Sci. U.S.A. 104, 7981–7986. (doi:10.1073/pnas.0611553104)

12. Gillespie JJ et al. 2012 A Rickettsia Genome Overrun by Mobile Genetic Elements Provides Insight into the Acquisition of Genes Characteristic of an Obligate Intracellular Lifestyle. J Bacteriol 194, 376–394. (doi:10.1128/JB.06244-11)

13. Cordaux R, Pichon S, Ling A, Pérez P, Delaunay C, Vavre F, Bouchon D, Grève P. 2008 Intense Transpositional Activity of Insertion Sequences in an Ancient Obligate Endosymbiont. Mol Biol Evol 25, 1889–1896. (doi:10.1093/molbev/msn134)

14. Schmitz-Esser S, Penz T, Spang A, Horn M. 2011 A bacterial genome in transition - an exceptional enrichment of IS elements but lack of evidence for recent transposition in the symbiont Amoebophilus asiaticus. BMC Evol Biol 11, 270. (doi:10.1186/1471-2148-11-270)

15. Kampfraath AA, Klasson L, Anvar SY, Vossen RHAM, Roelofs D, Kraaijeveld K, Ellers J. 2019 Genome expansion of an obligate parthenogenesis-associated Wolbachia poses an exception to the symbiont reduction model. BMC Genomics 20, 106. (doi:10.1186/s12864-019-5492-9)

16. Regassa L B. 2006 Spiroplasmas: evolutionary relationships and biodiversity. Front Biosci 11, 2983. (doi:10.2741/2027)

17. Hayashi M, Watanabe M, Yukuhiro F, Nomura M, Kageyama D. 2016 A Nightmare for Males? A Maternally Transmitted Male-Killing Bacterium and Strong Female Bias in a Green Lacewing Population. PLoS ONE 11, e0155794. (doi:10.1371/journal.pone.0155794)

18. Weinert LA, Werren JH, Aebi A, Stone GN, Jiggins FM. 2009 Evolution and diversity of Rickettsiabacteria. BMC Biol 7, 6. (doi:10.1186/1741-7007-7-6)

19. Cheng H, Concepcion GT, Feng X, Zhang H, Li H. 2021 Haplotype-resolved de novo assembly using phased assembly graphs with hifiasm. Nat Methods 18, 170–175. (doi:10.1038/s41592-020-01056-5)

20. Vaser R, Sović I, Nagarajan N, šikić M. 2017 Fast and accurate de novo genome assembly from long uncorrected reads. Genome Res. 27, 737–746. (doi:10.1101/gr.214270.116)

21. Laetsch DR, Blaxter ML. 2017 BlobTools: Interrogation of genome assemblies. F1000Res 6, 1287. (doi:10.12688/f1000research.12232.1)

22. Li H. 2018 Minimap2: pairwise alignment for nucleotide sequences. Bioinformatics 34, 3094–3100. (doi:10.1093/bioinformatics/bty191)

23. Camacho C, Coulouris G, Avagyan V, Ma N, Papadopoulos J, Bealer K, Madden TL. 2009 BLAST+: architecture and applications. BMC Bioinformatics 10, 421. (doi:10.1186/1471-2105-10-421)

24. Mikheenko A, Prjibelski A, Saveliev V, Antipov D, Gurevich A. 2018 Versatile genome assembly evaluation with QUAST-LG. Bioinformatics 34, i142–i150. (doi:10.1093/bioinformatics/bty266)

25. Manni M, Berkeley MR, Seppey M, Simão FA, Zdobnov EM. 2021 BUSCO Update: Novel and Streamlined Workflows along with Broader and Deeper Phylogenetic Coverage for Scoring of Eukaryotic, Prokaryotic, and Viral Genomes. Molecular Biology and Evolution 38, 4647–4654. (doi:10.1093/molbev/msab199)

26. Chklovski A, Parks DH, Woodcroft BJ, Tyson GW. 2023 CheckM2: a rapid, scalable and accurate tool for assessing microbial genome quality using machine learning. Nat Methods 20, 1203–1212. (doi:10.1038/s41592-023-01940-w)

27. Castelli M, Nardi T, Gammuto L, Bellinzona G, Sabaneyeva E, Potekhin A, Serra V, Petroni G, Sassera D. 2024 Host association and intracellularity evolved multiple times independently in the Rickettsiales. Nat Commun 15, 1093. (doi:10.1038/s41467-024-45351-7)

28. R Core Team. 2020 R: A language and environment for statistical computing.

29. Wickham, Hadley. 2016 ggplot2: Elegant Graphics for Data Analysis.

30. Wickham, H, Francois, R, Henry, L, Muller, K, Vaughan, D. 2023 dplyr: A Grammar of Data Manipulation.

31. Wickham, Hadley. 2023 forcats: Tools for Working with Categorical Variables (Factors).

32. Schwengers O, Jelonek L, Dieckmann MA, Beyvers S, Blom J, Goesmann A. 2021 Bakta: rapid and standardized annotation of bacterial genomes via alignment-free sequence identification: Find out more about Bakta, the motivation, challenges and applications, here. Microbial Genomics 7. (doi:10.1099/mgen.0.000685)

33. Eren AM et al. 2020 Community-led, integrated, reproducible multi-omics with anvi’o. Nat Microbiol 6, 3–6. (doi:10.1038/s41564-020-00834-3)

34. Tanizawa Y, Fujisawa T, Nakamura Y. 2018 DFAST: a flexible prokaryotic genome annotation pipeline for faster genome publication. Bioinformatics 34, 1037–1039. (doi:10.1093/bioinformatics/btx713)

35. Marçais G, Delcher AL, Phillippy AM, Coston R, Salzberg SL, Zimin A. 2018 MUMmer4: A fast and versatile genome alignment system. PLoS Comput Biol 14, e1005944. (doi:10.1371/journal.pcbi.1005944)

36. Wishart DS, Han S, Saha S, Oler E, Peters H, Grant JR, Stothard P, Gautam V. 2023 PHASTEST: faster than PHASTER, better than PHAST. Nucleic Acids Research 51, W443–W450. (doi:10.1093/nar/gkad382)

37. Xie Z, Tang H. 2017 ISEScan: automated identification of insertion sequence elements in prokaryotic genomes. Bioinformatics 33, 3340–3347. (doi:10.1093/bioinformatics/btx433)

38. Benson G. 1999 Tandem repeats finder: a program to analyze DNA sequences. Nucleic Acids Research 27, 573–580. (doi:10.1093/nar/27.2.573)

39. Couvin D et al. 2018 CRISPRCasFinder, an update of CRISRFinder, includes a portable version, enhanced performance and integrates search for Cas proteins. Nucleic Acids Research 46, W246–W251. (doi:10.1093/nar/gky425)

40. Kalyaanamoorthy S, Minh BQ, Wong TKF, Von Haeseler A, Jermiin LS. 2017 ModelFinder: fast model selection for accurate phylogenetic estimates. Nat Methods 14, 587–589. (doi:10.1038/nmeth.4285)

41. Minh BQ, Schmidt HA, Chernomor O, Schrempf D, Woodhams MD, Von Haeseler A, Lanfear R. 2020 IQ-TREE 2: New Models and Efficient Methods for Phylogenetic Inference in the Genomic Era. Molecular Biology and Evolution 37, 1530–1534. (doi:10.1093/molbev/msaa015)

42. Stecher G, Tamura K, Kumar S. 2020 Molecular Evolutionary Genetics Analysis (MEGA) for macOS. Molecular Biology and Evolution 37, 1237–1239. (doi:10.1093/molbev/msz312)

43. Halter T, Köstlbacher S, Rattei T, Hendrickx F, Manzano A, Horn M. In press. One to host them all: genomics of the diverse bacterial endosymbionts of the spider Oedothorax gibbosus. Microbial Genomics

44. El Karkouri K, Ghigo E, Raoult D, Fournier P-E. 2022 Genomic evolution and adaptation of arthropod-associated Rickettsia. Sci Rep 12, 3807. (doi:10.1038/s41598-022-07725-z)

45. Bordenstein SR, Reznikoff WS. 2005 Mobile DNA in obligate intracellular bacteria. Nat Rev Microbiol 3, 688–699. (doi:10.1038/nrmicro1233)

46. Wu Y, Aandahl RZ, Tanaka MM. 2015 Dynamics of bacterial insertion sequences: can transposition bursts help the elements persist? BMC Evol Biol 15, 288. (doi:10.1186/s12862-015-0560-5)

47. Huang D, Xia R, Chen C, Liao J, Chen L, Wang D, Alvarez PJJ, Yu P. 2024 Adaptive strategies and ecological roles of phages in habitats under physicochemical stress. Trends in Microbiology 32, 902–916. (doi:10.1016/j.tim.2024.02.002)

48. Bolaños LM, Servín-Garcidueñas LE, Martínez-Romero E. 2015 Arthropod–Spiroplasma relationship in the genomic era. FEMS Microbiology Ecology 91, 1–8. (doi:10.1093/femsec/fiu008)

49. Saillard C et al. 1987 Spiroplasma phoeniceum sp. nov., a New Plant-Pathogenic Species from Syria. International Journal of Systematic Bacteriology 37, 106–115. (doi:10.1099/00207713-37-2-106)

50. Philip RN, Casper EA, Anacker RL, Cory J, Hayes SF, Burgdorfer W, Yunker CE. 1983 Rickettsia bellii sp. nov.: a Tick-Borne Rickettsia, Widely Distributed in the United States, That Is Distinct from the Spotted Fever and Typhus Biogroups. International Journal of Systematic Bacteriology 33, 94–106. (doi:10.1099/00207713-33-1-94)

51. Lawson ET, Mousseau TA, Klaper R, Hunter MD, Werren JH. 2001 Rickettsia associated with male-killing in a buprestid beetle. Heredity 86, 497–505. (doi:10.1046/j.1365-2540.2001.00848.x)

