## Supplemental Figures S1-8 for "Concerted genome expansion of heritable symbionts in an insect host"

Supplementary Data

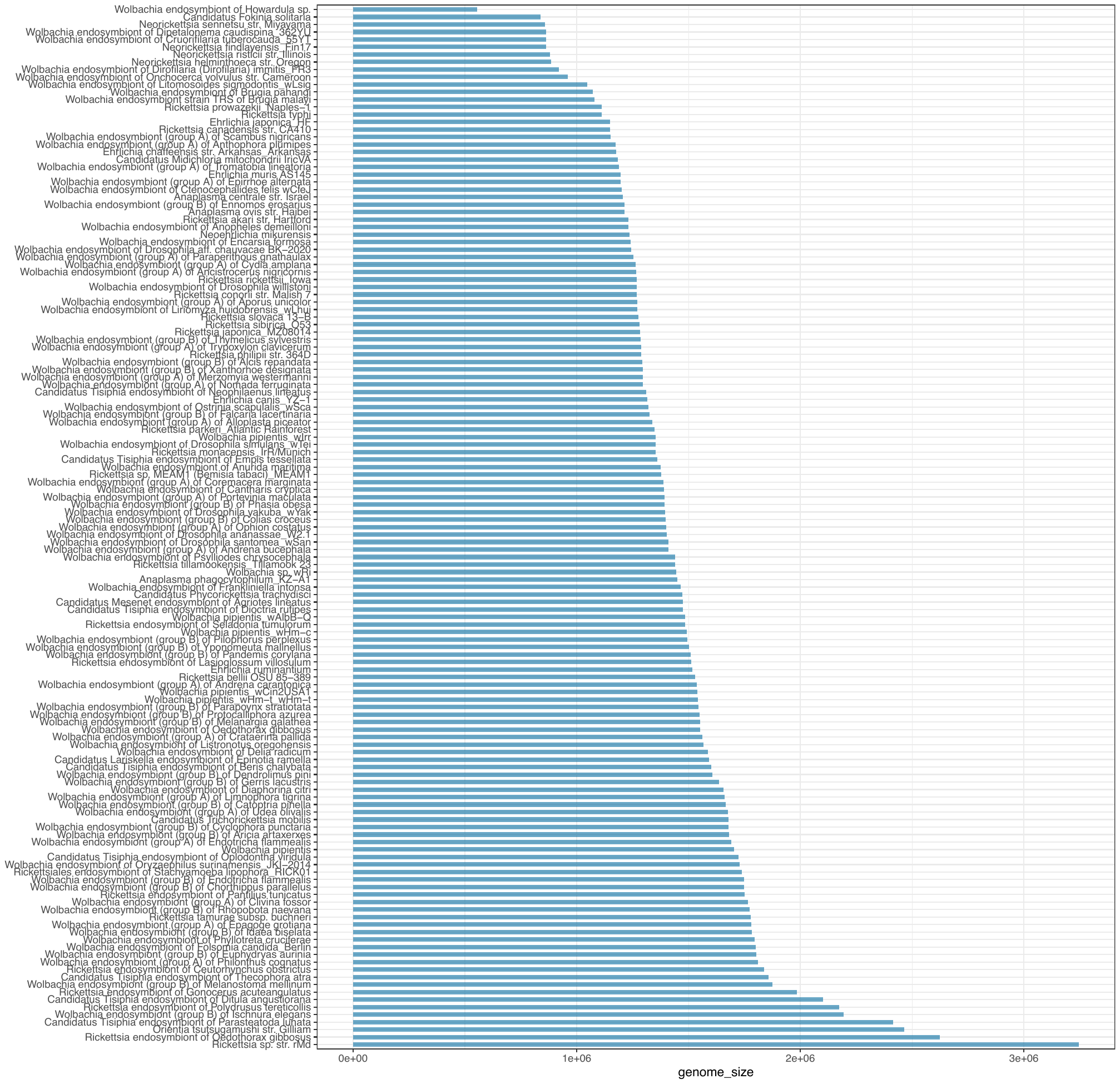

**Figure S1.** Genome sizes of sequenced complete Rickettsiales bacteria

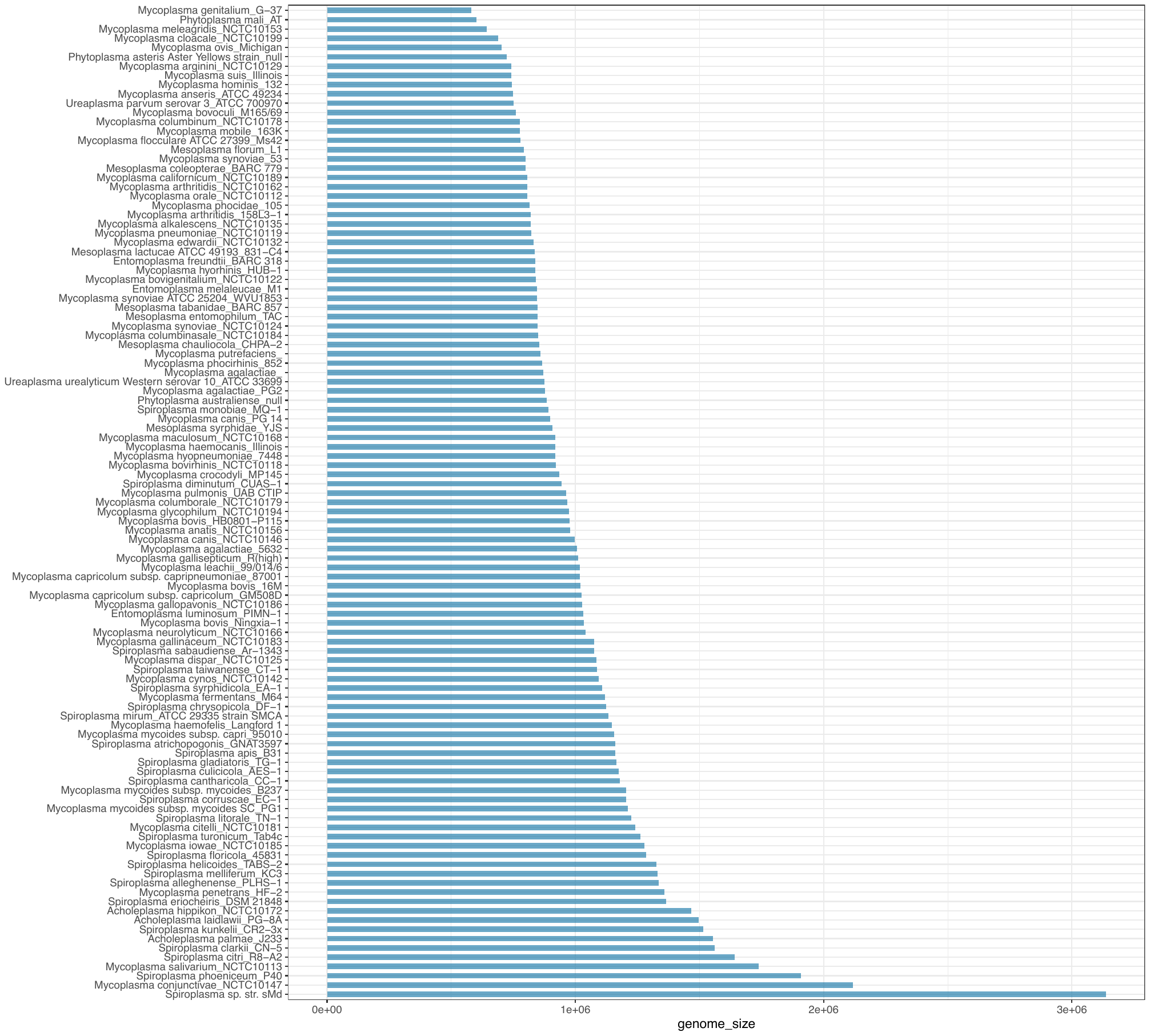

**Figure S2.** Genome sizes of sequenced complete Mollicute bacteria

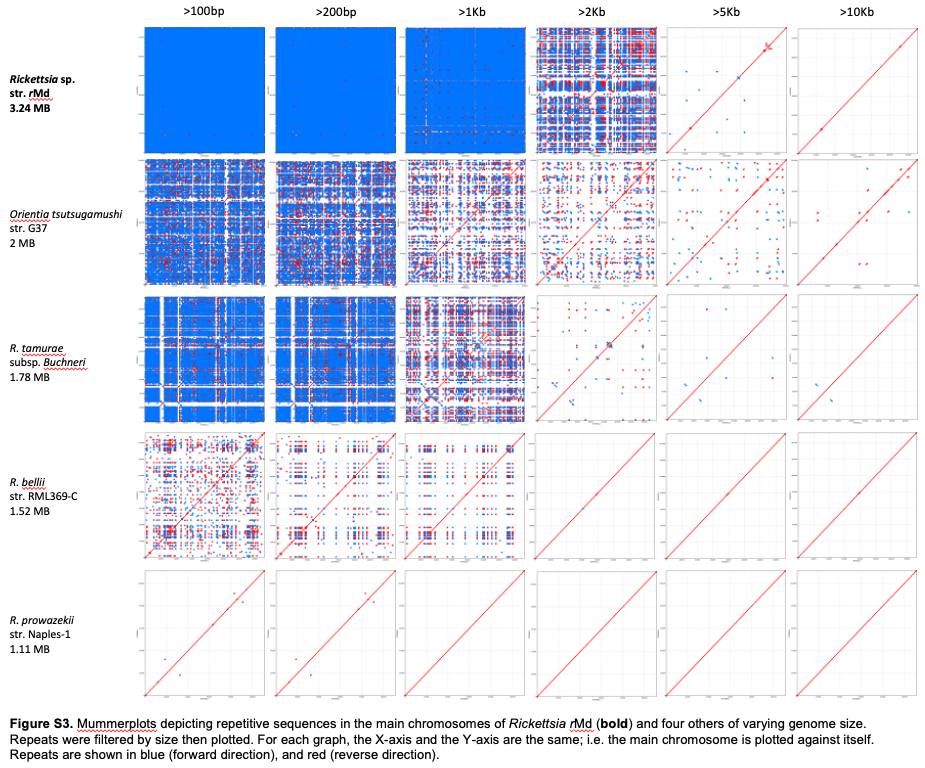

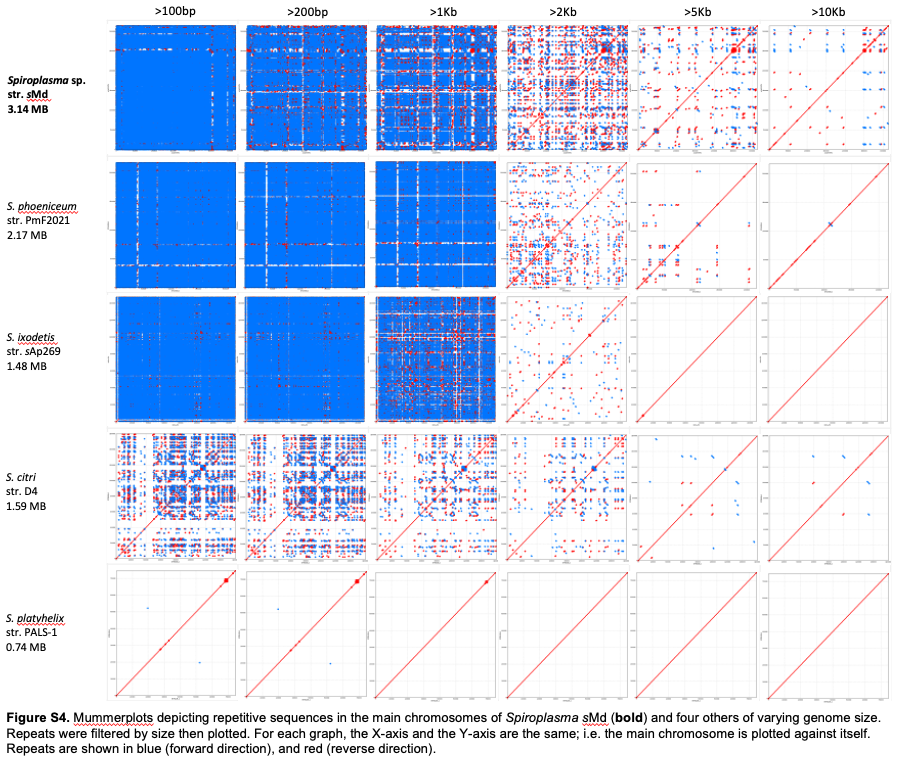

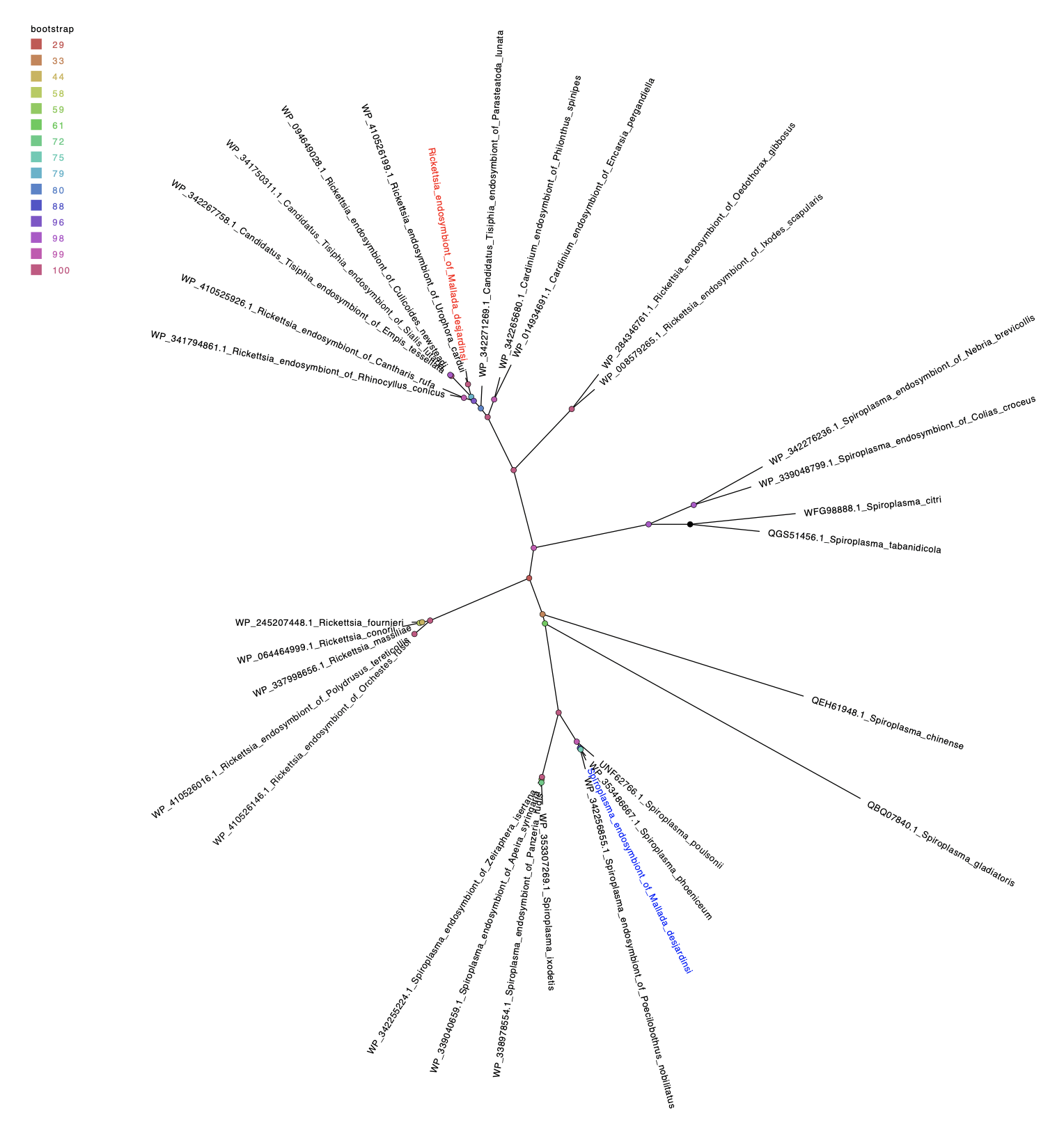

**Figure S5**. Star phylogeny (best-fit protein model according to Bayesian Information Criterion - VT+F+G4) of IS3 family transposase amino acid sequences from *Spiroplasma s*Md (in blue), *Rickettsia r*MD (in red) alongside selected strains from both *Rickettsia* and *Spiroplasma*.

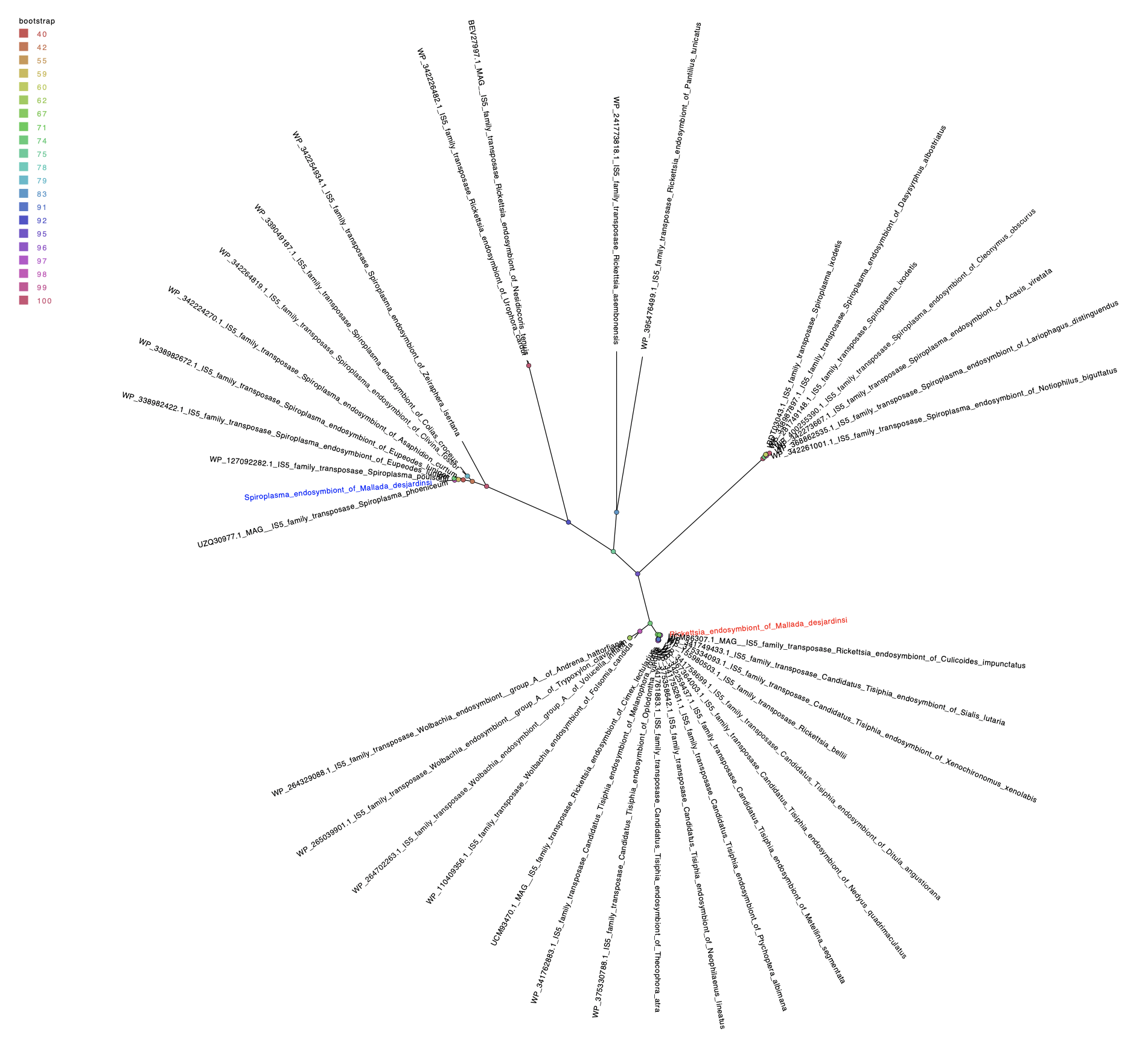

**Figure S6**. Star phylogeny (best-fit protein model according to Bayesian Information Criterion - JTT+F+G4) of IS5 family transposase amino acid sequences from *Spiroplasma s*Md (in blue), *Rickettsia r*MD (in red) alongside selected strains from both *Rickettsia* and *Spiroplasma*.

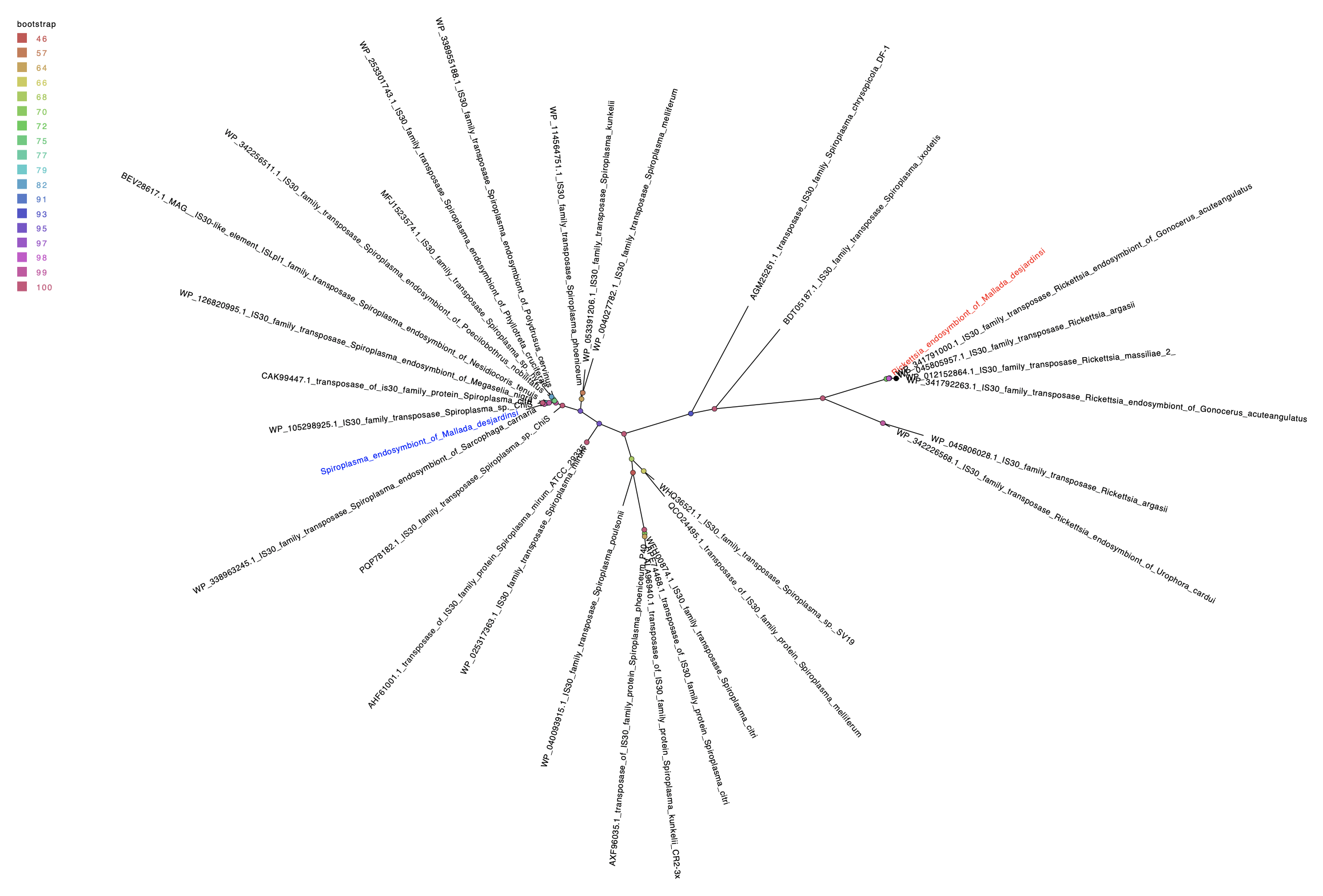

**Figure S7**. Star phylogeny (best-fit protein model according to Bayesian Information Criterion - VT+F+I+G4) of IS30 family transposase amino acid sequences from *Spiroplasma s*Md (in blue), *Rickettsia r*MD (in red) alongside selected strains from both *Rickettsia* and *Spiroplasma*.

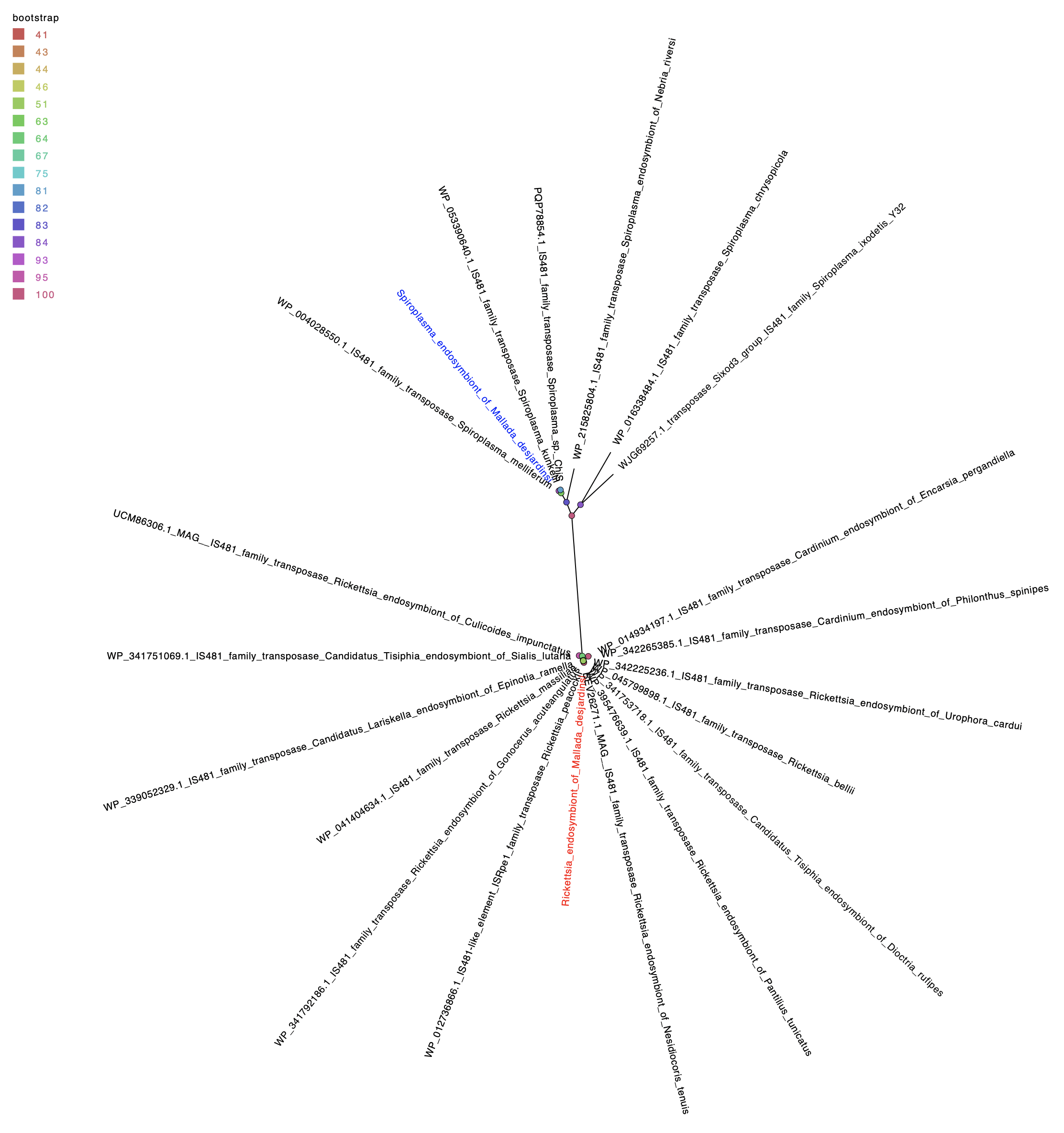

**Figure S8**. Star phylogeny (best-fit protein model according to Bayesian Information Criterion - Q.pfam+F+R3) of IS481 family transposase amino acid sequences from *Spiroplasma s*Md (in blue), *Rickettsia r*MD (in red) alongside selected strains from both *Rickettsia* and *Spiroplasma*.
