## Supplemental Figures S9-13 for "Concerted genome expansion of heritable symbionts in an insect host"

**Figure S9** *Rickettsia* kegg module analysis

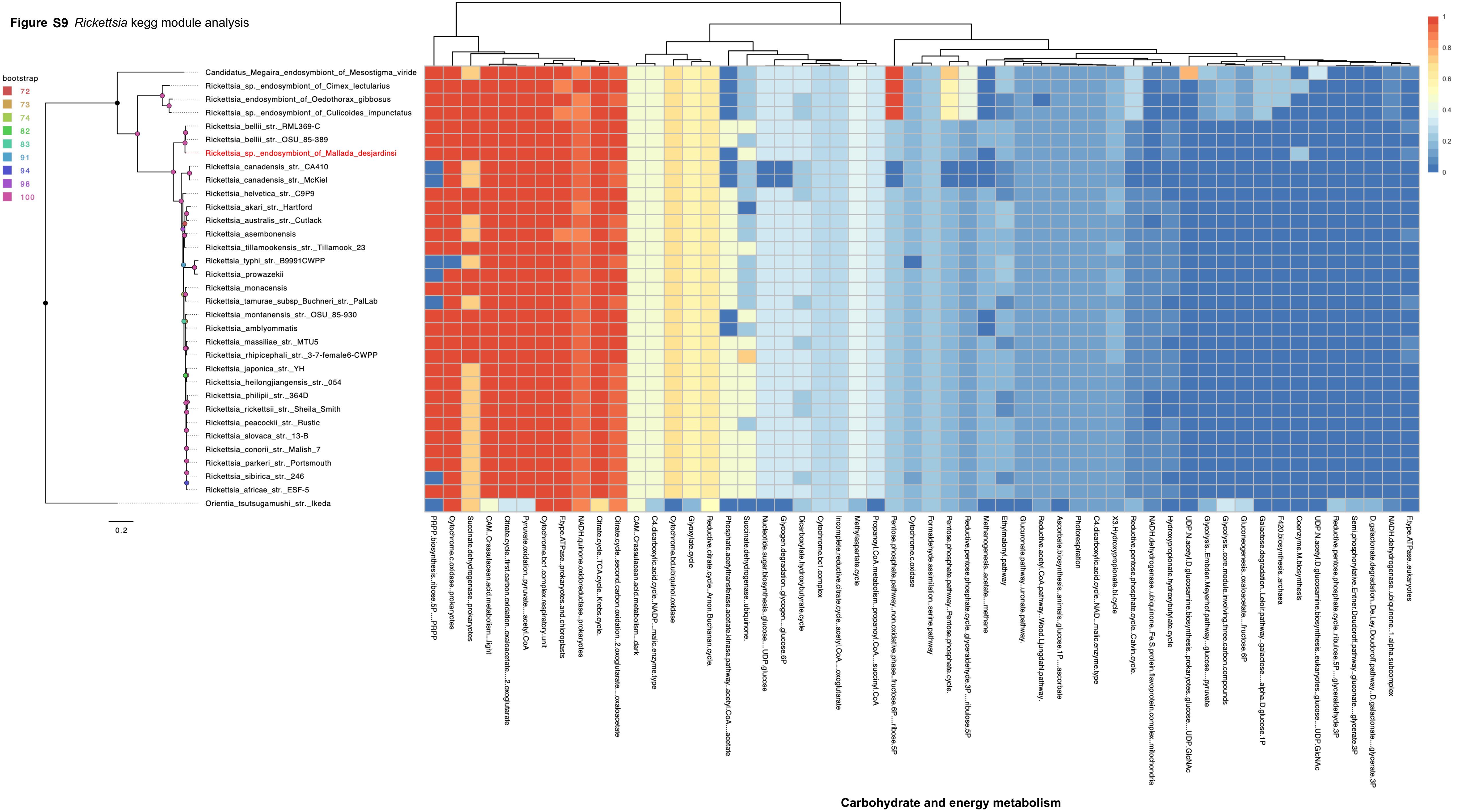

**Figure S10** *Rickettsia* kegg module analysis

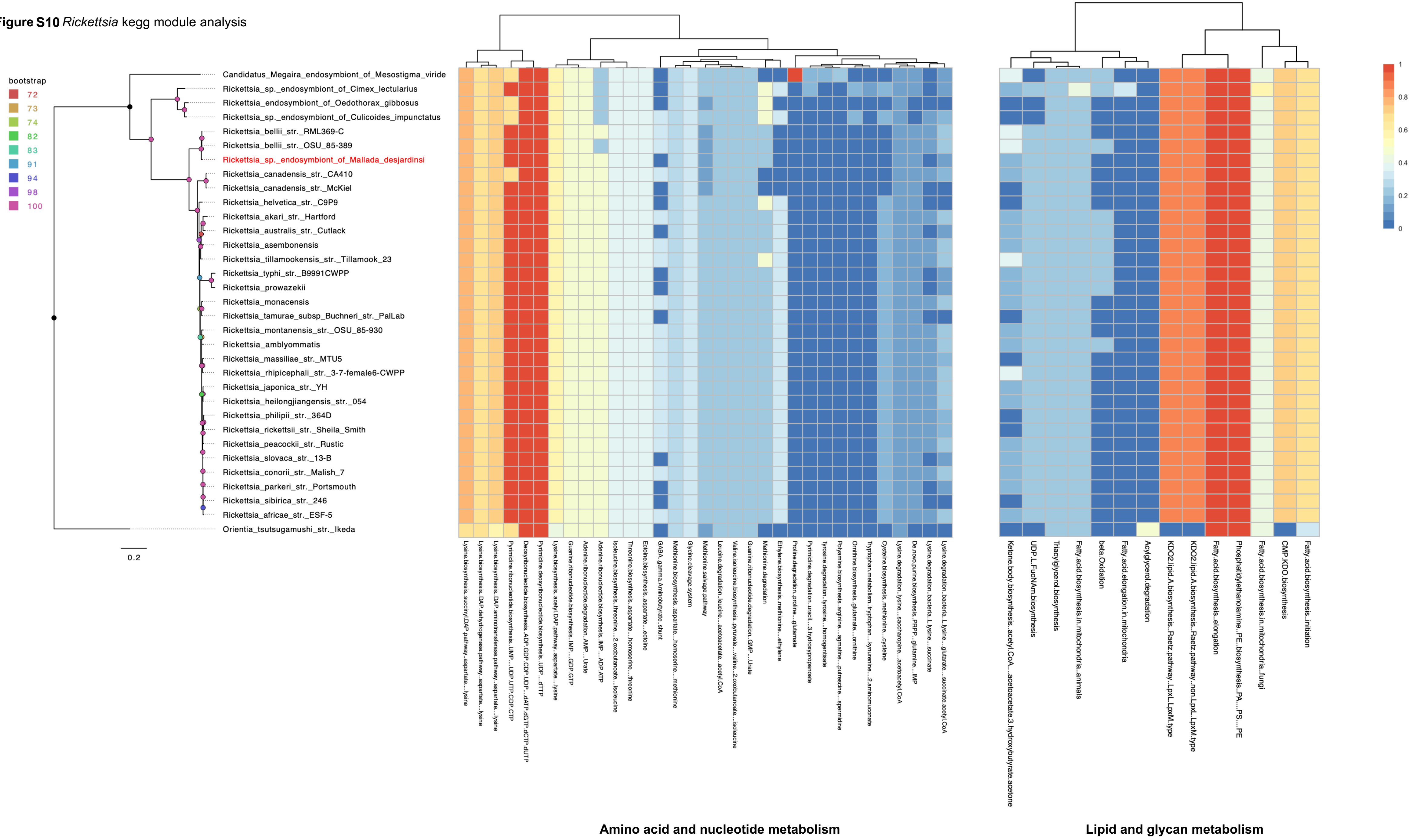

**Figure S11** *Rickettsia* kegg module analysis

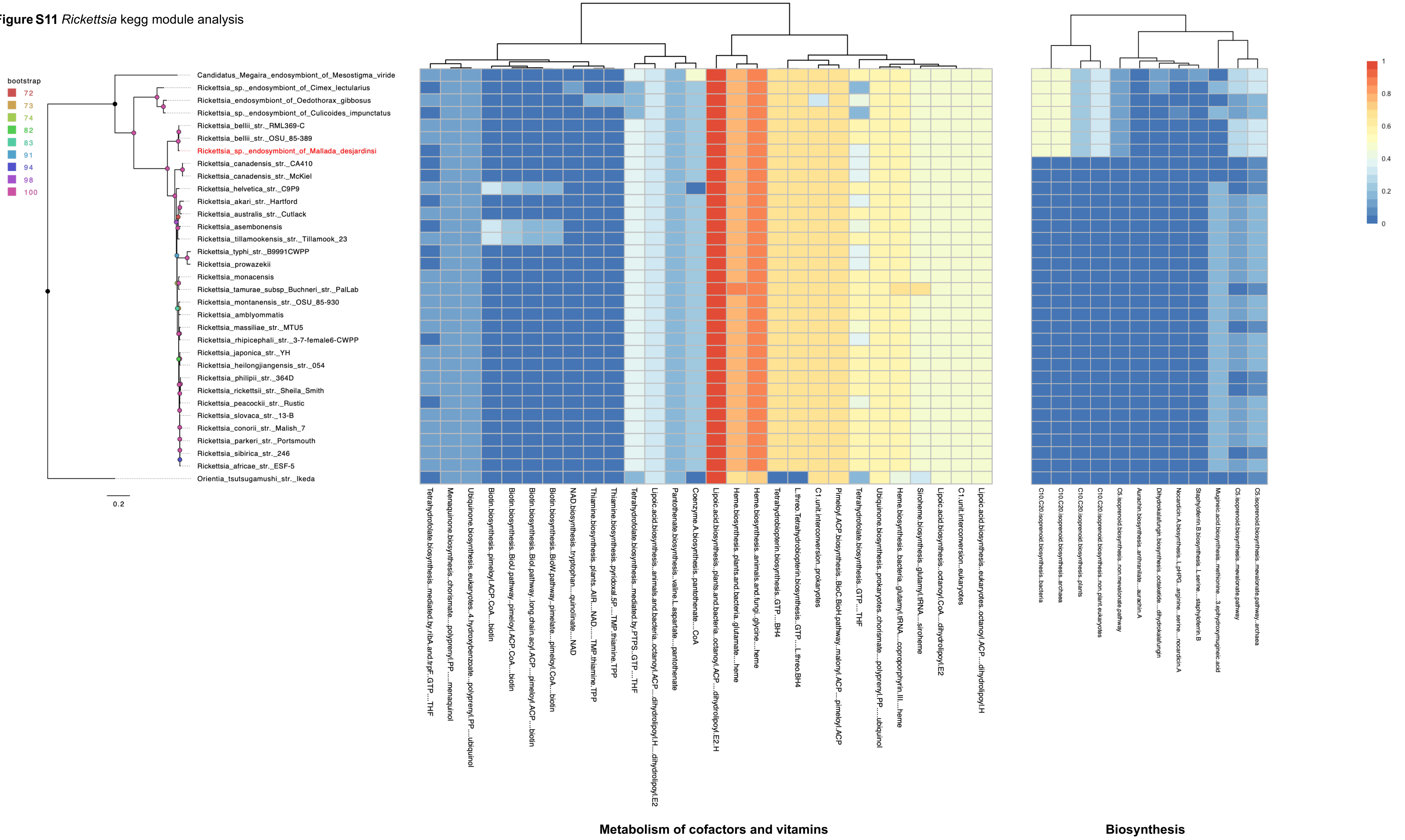

**Figure S12** *Spiroplasma* kegg module analysis

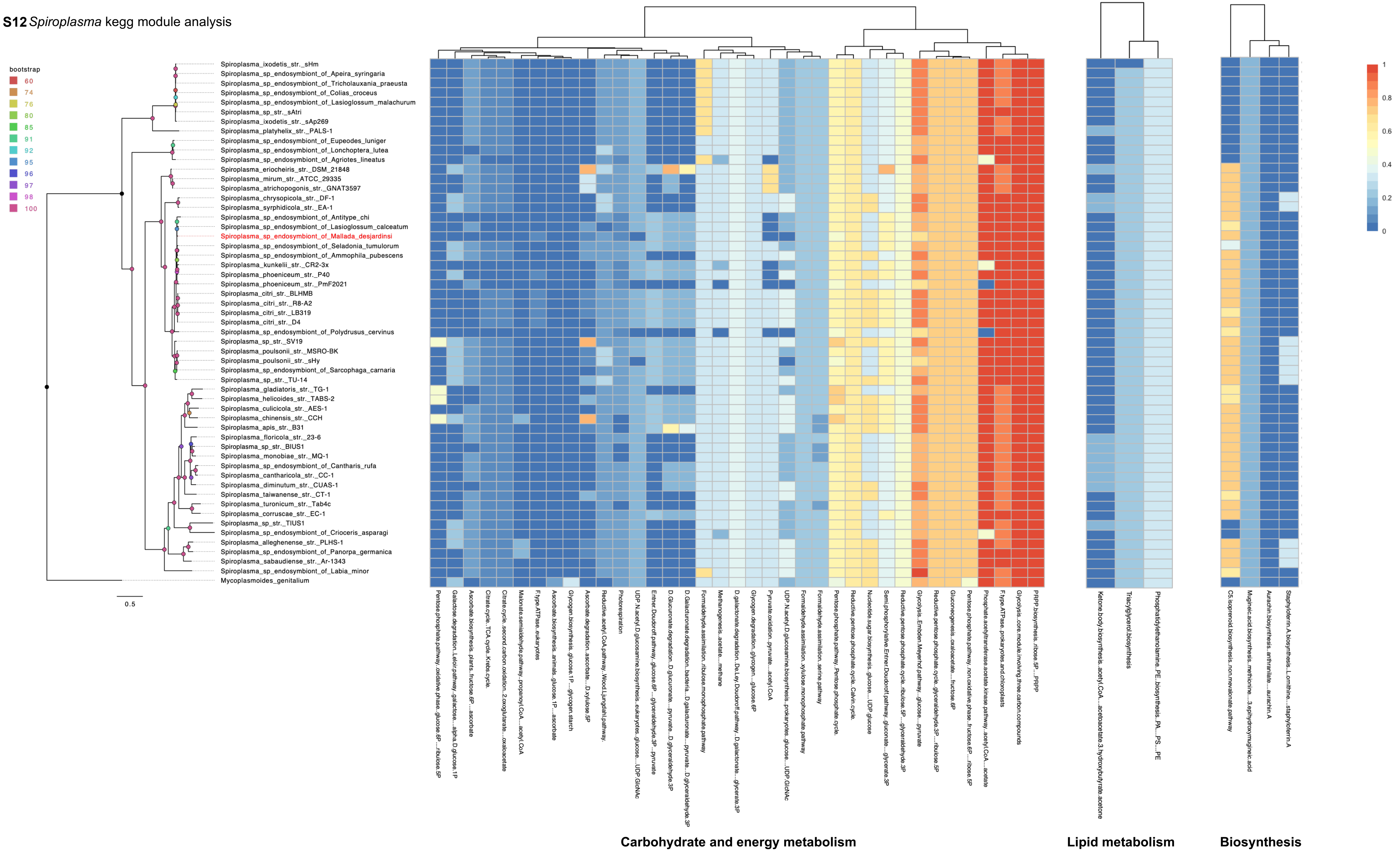

**Figure S13** *Spiroplasma* kegg module analysis

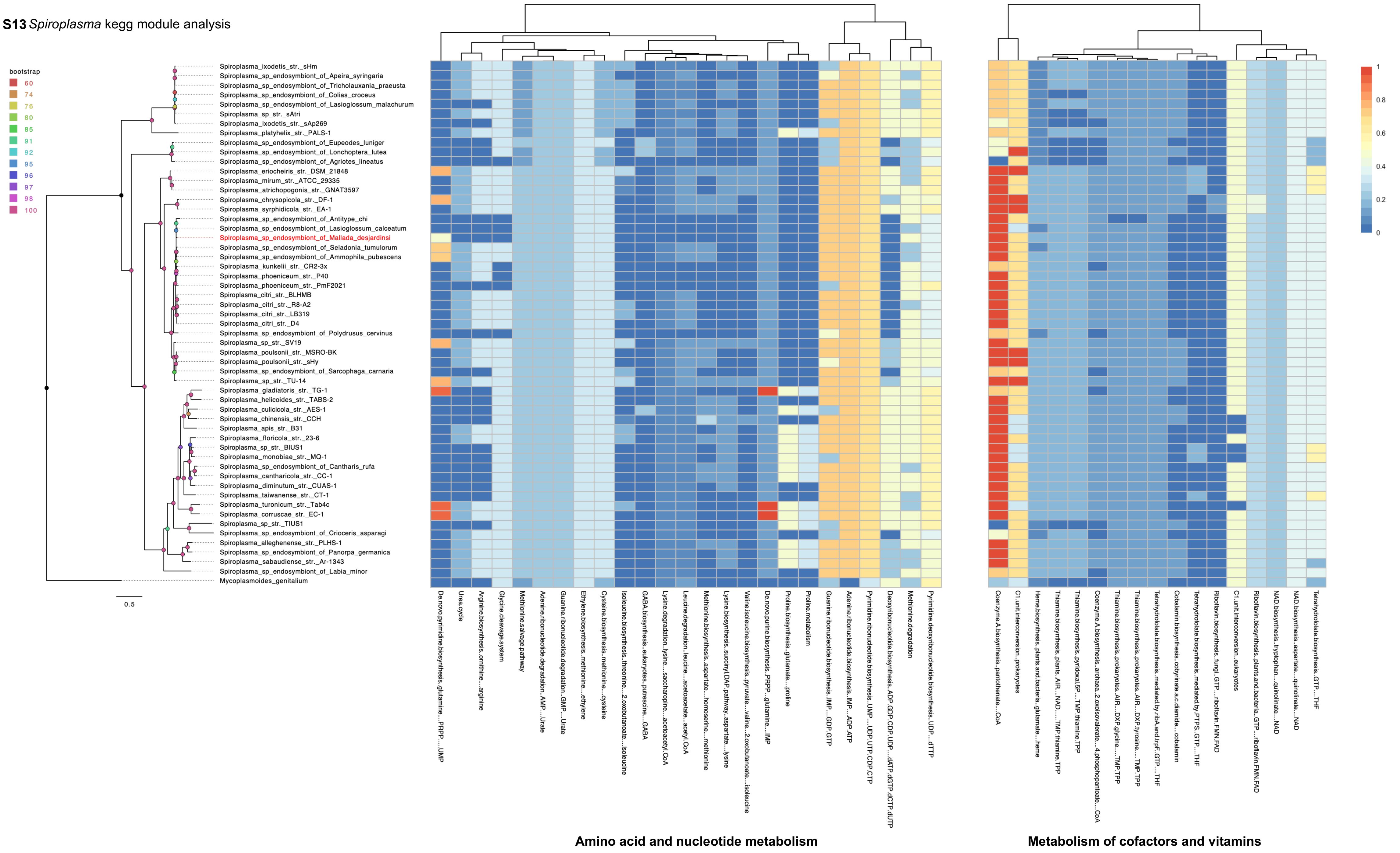
